# Unspecific expression in limited excitatory cell populations in interneuron-targeting Cre-driver lines can have large functional effects

**DOI:** 10.1101/848655

**Authors:** Daniel Müller-Komorowska, Thoralf Opitz, Shehabeldin Elzoheiry, Michaela Schweizer, Heinz Beck

## Abstract

Transgenic Cre-recombinase expressing mouse lines are widely used to express fluorescent proteins and opto-/chemogenetic actuators, making them a cornerstone of modern neuroscience. Particularly, the investigation of interneurons has benefitted from the ability to target genetic constructs to defined cell types. However, the cell type specificity of some mouse lines has been called into questions. Here we show for the first time the functional consequences of unspecific expression in a somatostatin-Cre (SST-Cre) mouse line. We find large optogenetically evoked excitatory currents originating from unspecifically targeted CA3 pyramidal cells. We also used public Allen Brain Institute data to estimate expression specificity in other Cre lines. Another SST-Cre mouse lines shows comparable unspecificity, whereas a Parvalbumin-Cre mouse line shows much less unspecific expression. Finally, we make suggestions to ensure that the results from in-vivo use of Cre mouse lines are interpretable.

## 2 Introduction

Transgenic Cre-recombinase expressing mouse lines are widely used in modern neuroscience to specifically direct the expression of fluorescent proteins or opto- and chemogenetic actuators to neuronal subtypes. Accordingly, they are a key element of most neuronal perturbation studies. Cre driver mouse lines have been extensively used to examine the function of interneuron subtypes in-vitro and in-vivo, with increasing numbers of Cre mouse lines for specific molecular markers of different interneuron subtypes (Taniguchi et al., 2011). Very commonly used are mice expressing Cre in subsets of GABAergic interneurons under the parvalbumin (PV) or somatostatin (SST) promoters. Those lines have allowed to target two main categories of interneurons. In the hippocampus, PV^+^ cells include fast spiking basket cells, axo-axonic cells, and interneuron types targeting proximal dendrites of pyramidal cells. SST^+^ cells, on the other hand, are regularly spiking and inhibit pyramidal cells at their distal dendrites (Lovett-Barron et al., 2012; Pelkey et al., 2017).

Commonly used Cre-lines have been widely assumed to be specific, with Cre-expression confined to the cells of interest. However, this assumption has been called into question in some cases. For example, in the widely used somatostatin–IRES-Cre line (Taniguchi et al., 2011), a population of 5% of Cre-reporter positive cells were found to be fast-spiking PV^+^ cells (Hu, Cavendish, & Agmon, 2013; Mikulovic, Restrepo, Hilscher, Kullander, & Leão, 2015). In the hippocampal CA1 subfield, this mouse line also targets a small (6%) population of fast-spiking interneurons (Hu et al., 2013; Mikulovic et al., 2015). Particularly opto- and chemogenetic studies often depend on highly specific expression patterns to disseminate the function of neuronal subtypes. Even though these findings are worrisome, one defense of such mouse lines is that the absolute number of unspecifically targeted cells is small. One could therefore assume that the observed in-vitro and in-vivo effects are dominated by the interneuron type in question.

Here we show that in SST-Cre mice (Savanthrapadian et al., 2014), recombination is not only induced in GABAergic interneuron types. In addition, recombination also occurs in a small subset of excitatory neurons largely confined to the CA3 pyramidal cell layer. Moreover, we find powerful functional effects of optogenetic activation that are not only contaminated by unspecifically expressing glutamatergic cells but are completely lacking any interneuron contribution. Finally, we were also unable to find anatomical or functional differences between unspecifically targeted cells and canonical CA3 pyramidal cells. This suggests that these cells are not a specific subtype of CA3 pyramidal cell. Further control experiments should be carried out in a region-specific manner, prior to the use of Cre-lines for the investigation of circuit function in behavior.

## 3 Methods

### 3.1 Transgenic Animals

All animal experiments were carried out according to the guidelines stated in Directive 2010/63/EU of the European Parliament on the protection of animals used for scientific purposes and were approved by authorities in Nordrhein-Westfalen (Landesamt für Natur, Umwelt und Verbraucherschutz Nordrhein Westfalen (LANUV), AZ 84-02.04.2014.A254).

The SST-Cre mouse line was kindly provided to us by Marlene Bartos and was described previously (Savanthrapadian et al., 2014). It was also used in a more recent study (Eyre & Bartos, 2019). Animals were bred hemizygous and were genotyped for Cre recombinase using the forward primer CCATCTGCCACCAGCCAG and the reverse primer TCGCCATCTTCCAGCAGG. Animals with an amplified fragment at 281bp were classified as transgenic. For the cross-breeding experiments (**Figure 1C**) we used the Ai14 reporter line (Jackson Laboratories Stock No 007914).

**Figure 1:**
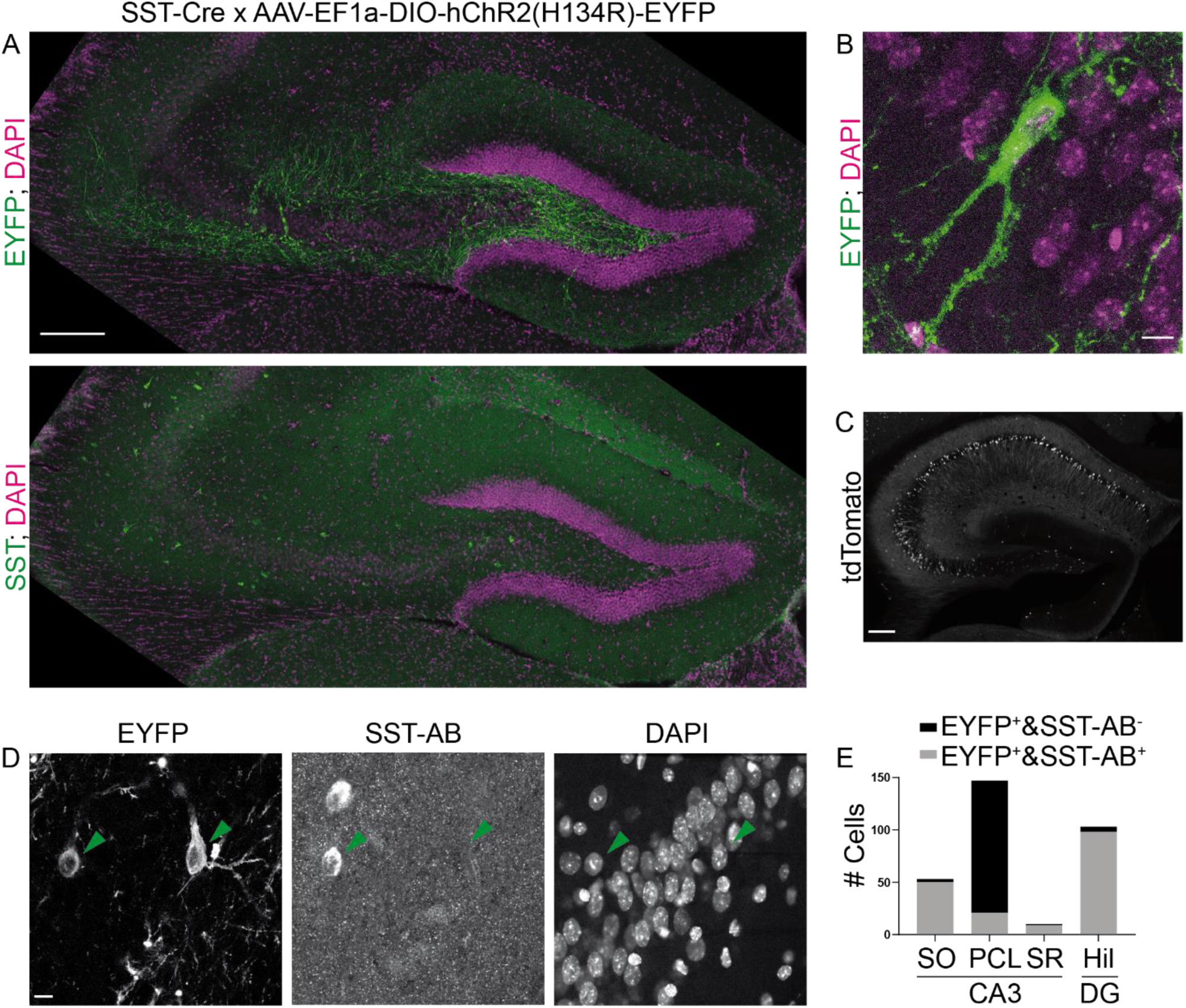
The SST-Cre line is not specific for SST^+^ interneurons in CA3. A) CA3 and the hilus of the dentate gyrus were virally transduced by intracranial stereotactic injection with a Cre dependent, EYFP expressing construct. Lower Image shows SST staining in the same slice. 10x objective. Scalebar 200µm. Contrast adjusted for visualization. B) EYFP positive cell in CA3 PCL and two apical dendrites in stratum lucidum. Scalebar 10µm. C) Hippocampal tdTomato signal in the SST-Cre mouse line crossed with at tdTomato reporter line. 10x magnification. Scalebar 200µm. D) Example images showing 2 EYFP^+^ cells, one in SO and one in the PCL. The cell in SO is also SST positive, the cell in the PCL is SST negative. 40x magnification. Scalebar 10µm. E) Quantification of SST colocalization for 40x images.

### 3.2 Stereotaxic intracranial viral injections

Animals were anesthetized with a ketamine/rompun or a fentanyl/midazolam/medetomidin mixture i.p. Animals also received ketoprofen analgesia (5 mg/kg, 0.1 ml/10 g body weight) before the surgery and daily 2 days after the surgery. Viral particles (250 nl at a rate of 100 nl/min) were injected into CA3/hilus of the right hemisphere at the following coordinates relative to bregma: 2.3 mm posterior; 1.6 mm lateral; 2.5 mm ventral. We used rAAV-Ef1a-DIO-hChR2(H134R)-EYFP-WPRE-pA (Received as a gift from Karl Deisseroth, Addgene plasmid # 20298; http://n2t.net/addgene:20298; RRID:Addgene_20298) for Cre-mediated opsin expression, AAV1/2-Ef1a-DIO-Syp-miniSOG-t2A-mCherry-WPRE-hpA (Received as a gift from Roger Tsien; Shu et al., 2011) for electron microscopy experiments and AAV1/2.Syn-hChR2(H134R)-EYFP (Received as a gift from Karl Deisseroth, Addgene plasmid # 26973; http://n2t.net/addgene:26973; RRID:Addgene_26973) for general expression. Cholera Toxin subunit B (CT-B), Alexa Fluor 555 conjugate (C-34775, Thermo Fischer) was injected into CA1 at bregma coordinates: 1.9 mm posterior; 1.5mm lateral; 1.7 ventral. Mice were used for electrophysiological experiments 4 to 5 weeks after viral injection.

### 3.3 Somatostatin immunostaining

Animals were transcardially perfused with 4% PFA and the brains were post-fixed with 4% PFA overnight at 4°C. The brains were washed in PBS the next day and slices of the dorsal hippocampus were cut on a vibratome (HM 650V; Thermo Scientific) at 50µm. After washing, slices were left in a blocking solution, consisting of 3% BSA in 0.25% PBS-T, for 2h at room temperature (RT). Then the primary antibody, rabbit anti-SST (T-4102, Peninsula Laboratories International), was applied 1:500 in blocking solution overnight shaking at 4°C. The following day slices were left at RT for 30 minutes and washed in blocking solution. The secondary antibody, donkey anti-rabbit IgG, Alexa fluor 647 (ab150075, Abcam), was applied 1:500 overnight shaking at 4°C. Finally, slices were washed, stained with 1:1000 DAPI for 30min at RT shaking and mounted with aqua-polymount. The SST staining for the choleratoxin-B (CT-B) injected animals followed a slightly different protocol where slices were blocked with 5% donkey serum instead of BSA and the secondary antibody was donkey anti-rabbit IgG FITC 1:500 (ab6798, Abcam). Colocalization was quantified manually from 40x images.

### 3.4 In-vitro electrophysiology

Adult mice were anesthetized with isofluorane, rapidly decapitated and the dissected brains were transferred to ice cold, carbogenated artificial cerebrospinal fluid with sucrose (ACSF; in mM: NaCl, 60; sucrose, 100; KCL, 2.5; NaH_2_PO_4_, 1.25; NaHCO_3_, 26; CaCl_2_, 1; MgCl_2_, 5; glucose, 20; from Sigma-Aldrich) and sliced to 300µm. Slices were then transferred to ACSF at 37°C and left for 20 minutes. They were then transferred to carbogenated ACSF without sucrose (NaCl, 125; KCL, 3.5; NaH_2_PO_4_, 1.25, NaHCO_3_, 26; CaCl_2_, 2; MgCl_2_, 2; glucose, 20; from Sigma-Aldrich) and were used for experiments after at least one hour. All experiments were performed in the same ACSF without sucrose at RT. The intracellular solution for voltage clamp experiments contained in mM: Cs methanesulfonate, 120; MgCl_2_, 0.5; 2-(4-(2-Hydroxyethyl)-1-piperazinyl)-ethansulfonsäure (HEPES), 5; Ethylenglycol-bis(aminoethylether)-N,N,N′,N′-tetraessigsäure (EGTA), 5; Adenosine 5′-triphosphate disodium salt (Na_2_-ATP), 5; N-(2,6-Dimethylphenylcarbamoylmethyl)triethylammonium chloride (QX 314 Cl^-^), 5; from Sigma Aldrich. For pharmacology we furthermore used 10µM gabazine (SR 95531 hydrobromide; Tocris), 1µM tetrodotoxin (TTX, Tocris), 200µM 4-aminopyridine (4-AP, Sigma Aldrich), 50µM 6-Cyano-7-nitroquinoxaline-2,3-dione disodium salt (CNQX, Tocris), 200µM D-(-)-2-Amino-5-phosphonopentanoic acid (D-AP5, Tocris). All these compounds were applied in the recording chamber for at least 10 minutes before continuing measurements. Most were applied for 20 minutes.

Patch clamp experiments were performed with an Axopatch 200B and digitized on a Digidata 1322A or Digidata 1550B plus HumSilencer (Molecular Devices). Light stimulation was performed with an Omicron Luxx 473nm laser attached to a light fiber submerged in the ACSF. Light stimuli were 5ms long unless otherwise stated.

For the conductance analysis we assumed a chloride reversal potential of −80mV (−78.9mV calculated with Nernst equation) and a cation reversal potential of 0mV. The excitatory conductance was calculated from a current trace measured at a holding voltage near the chloride reversal with gabazine washed-in, to ensure pure excitatory response. To isolate the inhibitory conductance, we subtracted the pure excitatory response at a depolarized holding voltage from the mixed response in normal ACSF.

In **Figure 2C** we only included cells that showed complete block by TTX wash-in. We excluded one cell that did not show complete block, which is likely due to a wash-in failure.

**Figure 2:**
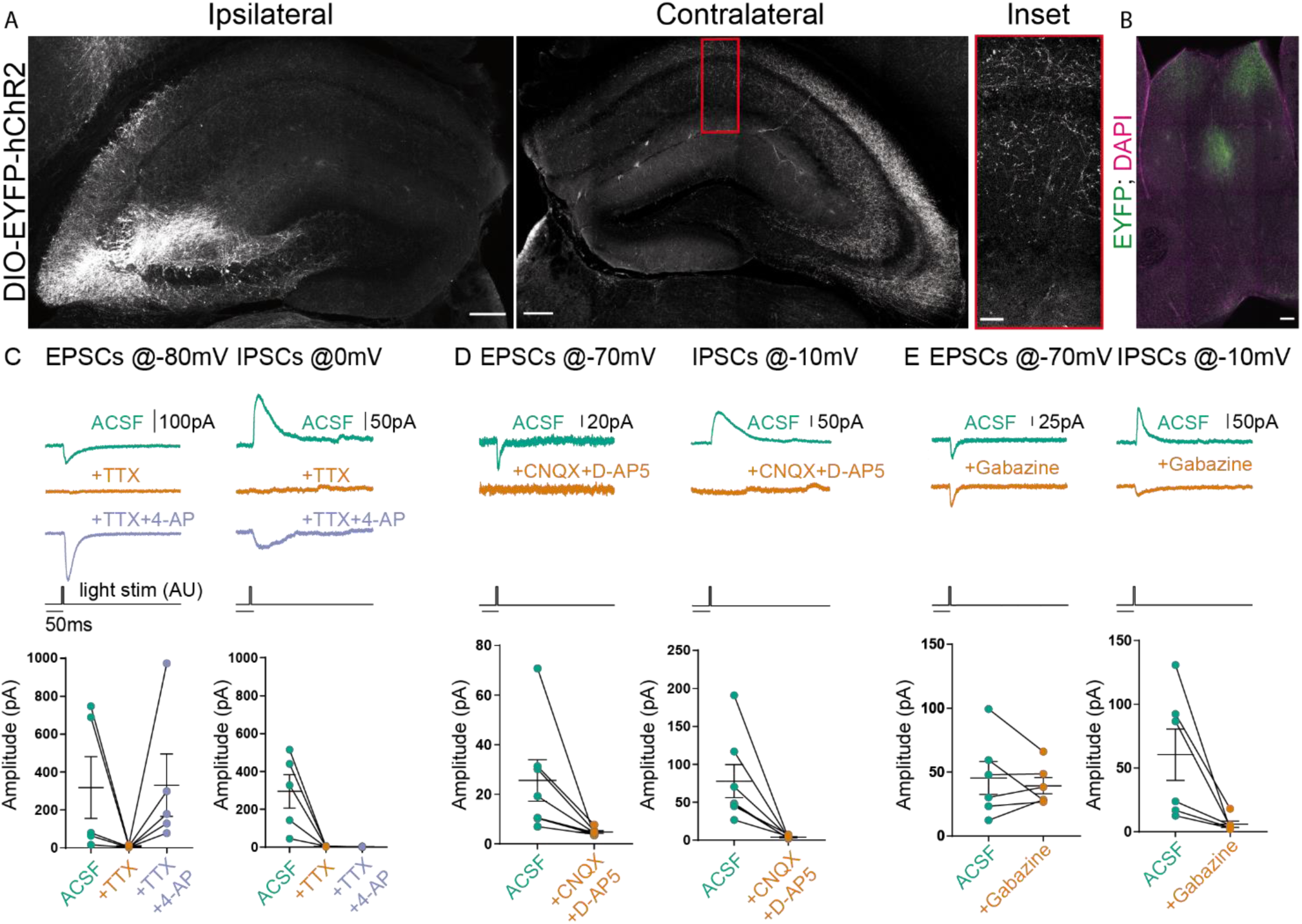
Stimulation of contralateral projections of CA4 neurons in the SST-Cre mouse line. A) Confocal images from post-fixed acute slices of the ipsilateral injection site (left) and the contralateral hippocampus (right). The inset shows fluorescent fiber signal in the contralateral hemisphere. Scalebars: 200µm. Inset: 100µm. B) Slice showing the projection from CA3 to the septum in the SST-Cre line. Scalebar 100µm. C), D), E) EPSCs and IPSCs measured in contralateral CA1 pyramidal cells. Light stimulus is 5ms long with 26mW total light-fiber output. C) Application of TTX alone abolished both excitatory and inhibitory currents. However, co-application of TTX+4-AP recovered EPSCs but not IPSCs (except in one cell). Ratio t-test of dependent samples between ACSF and +TTX+4-AP one-tailed: EPSCs, p = 0.1412, t=1.241, df=4; IPSCs, p < 0.0001, t=13.18, df =4. D) Application of CNQX+D-AP5 abolishes both EPSCs and IPSCs. Ratio t-test of dependent samples one-tailed: EPSCs, p=0.0017, t=4.681, df=6; IPSCs, p<0.0001, t=8.082, df=6. E) Application of Gabazine does not affect EPSCs but inhibits IPSCs. Ratio t-test of dependent samples one-tailed: EPSCs, p= 0.48186, t=0.04799, df=5; IPSCs, p= 0.0021, t=4.947, df=5. All Responses were recorded at 26mW fiber output.

### 3.5 Electron microscopy with miniSOG photooxidation

SST-Cre animals were virally transduced with EF1a-DIO-Syp-miniSOG-T2A-mCherry. Three weeks later, mice were transcardially perfused with Ringer solution followed by 4% formaldehyde in 0.15 M cacodylate-buffer. Brains were removed and post-fixed overnight at 4°C. Coronal slices (100µm) were taken on a vibratome and slices with distinct mCherry fluorescence were chosen. Slices were fixed with 2% Glutaraldehyde for 30 min, washed with ice-cold cacodylate-buffer, and blocked for 20 min in solution containing 20mM glycine, 10mM KCN, and 20mM aminotriazoline in cacodylate-buffer. For photooxidation, slices were immersed in freshly prepared and filtered (0.22um) 3,3’-diaminobenzidine (DAB) solution (1mg/ml DAB in 0.1 M cacodylate-Buffer, pH 7.4) that was aerated with oxygen. The miniSOG was activated with green light (FITC filterset: EX470/40, DM510, BA520) applied through a LUMPlanFl 60x NA 0.90 W at an inverted Olympus microscope equipped with a 100 W HBO-Lamp. Light was applied for 20 min and fresh DAB solution was exchanged after 10 min. After illumination, slices were stored in cacodylate-buffer for further processing.

After photoconversion the converted region containing DAB reaction product in the hippocampus was documented and images were taken at a Zeiss Axiophot light microscope. Thereafter the sections were rinsed three times in 0.1 M sodium cacodylate buffer (pH 7.2-7.4) (Sigma-Aldrich, Germany) and incubated with 1 % osmium tetroxide (Science Services, Germany) in cacodylate buffer for 20 minutes on ice. The osmication of sections was followed by dehydration through ascending ethyl alcohol concentration steps and rinsing twice in propylene oxide (Carl Roth, Germany). Infiltration of the embedding medium was performed by immersing the sections first in a mixture of 2:1 of propylene oxide and Epon (Carl Roth, Germany) then in a 1:1 mixture and finally in neat Epon and polymerised at 60°C for 48 hours. The region of interest was dissected and Ultrathin sections (60 nm) were prepared with a Leica Ultracut UC7. Images were taken using an EM902 transmission electron microscope (Zeiss, Germany) equipped with a CCD in lens 2K digital camera and running the ImageSP software (Tröndle, Moorenweis, Germany).

### 3.6 Quantification and Statistical Analysis

We used python with matplotlib (Hunter, 2007) and GraphPad Prism for plotting. Electrophysiological data were analysed manually in Clampfit (Molecular Devices) or with python and numpy (van der Walt, Colbert, & Varoquaux, 2011). To load .abf files into python we used the python-neo package (Garcia et al., 2014). GraphPad Prism was used for statistical analysis. We used the t-test to compare 2 groups and two-way ANOVA to compare two groups across multiple conditions.

For quantification of Allen Brain Institute data (Oh et al., 2014), we used the Allen Software Development Kit to download .jpg images. tdTomato positive cells were segmented by maximum entropy thresholding, erosion, dilation and the particle counter in ImageJ (Schindelin et al., 2012). Colocalization with fluorescent in-situ hybridization probe was assessed manually. In total we quantified 23 images of dorsal hippocampus from 4 experiments (Table 1). A detailed technical description can be found in the Transgenic Characterization whitepaper: http://help.brain-map.org/display/mouseconnectivity/Documentation

**Table 1:** Experiments and images from the Allen Brain Institute used for the quantification in **Figure 5**. All images can be found here: http://connectivity.brain-map.org/transgenic

| Line | Experiment | Img ID |  |  |  |  |  |
| --- | --- | --- | --- | --- | --- | --- | --- |
| SST-IRES-Cre | 182530118 | 182530130 | 182530134 | 182530136 | 182530140 | 182530142 | 182530156 |
|  | 167643437 | 167643500 | 167643502 | 167643504 | 167643514 | 167643516 |  |
| Pvalb-IRES-Cre | 81657984 | 81636703 | 81636705 | 81636709 | 81636711 | 81636713 | 81636715 |
|  | 111192541 | 111192610 | 111192612 | 111192625 | 111192627 | 111192629 |  |

## 4 Results

### 4.1 The SST-Cre line is not specific for SST^+^ interneurons in CA3

SST positive interneurons in CA3 are located predominantly in stratum oriens (SO) and stratum radiatum (SR). SST positive cells have a characteristic dendrite morphology, with most of the dendritic arbor confined to the same sublayer as the soma (Freund & Buzsáki, 1996). We expressed a construct that leads to Cre-dependent expression of EYFP in the CA3 region of heterozygous SST-Cre mice using rAAV-dependent gene transfer. We found EYFP expression in cells of the pyramidal cell layer (PCL; **Figure 1A**). In stratum oriens and stratum radiatum, cells also expressed EYFP but the signal there was almost dominated by the neuropil. EYFP^+^ cells in the PCL showed features typical for CA3 pyramidal cells (**Figure 1B**) such as thorny excrescences on apical dendrites.

To determine if these EYFP^+^ cells are also SST^+^, we immunostained for SST. This revealed that EYFP expression was highly specific for SST^+^ interneurons in SO, where 50/53 EYFP^+^ cells expressed SST. Similarly, in SR 9/10 EYFP^+^ cells expressed SST. In marked contrast we found that a minority of EYFP^+^ cells in the pyramidal cell layer of CA3 coexpressed SST (21/147 cells, **Figure 1D,E**). In addition to viral gene transfer of a reporter construct, we also crossed mice of our SST-Cre line with a tdTomato reporter mouse (line Ai14, see methods). As in the previous experiment, we found that most reporter positive cells were in the PCL and showed pyramidal like dendritic morphology in CA3, CA2 and CA1. Interestingly, we also found a very small number of granule cell-like neurons in the granule cell layer of the dentate gyrus (**Figure 1C**) that were not observed in virally transduced animals.

These results show that Cre recombinase is not only targeted to SST^+^ interneurons in the hippocampus. It is also expressed in pyramidal-like neurons within the pyramidal cell layer that are devoid of detectable somatostatin levels. In marked contrast, the SST-Cre mouse line showed high local specificity in CA3 SO, SR and the hilus of the dentate gyrus.

### 4.2 Commissural projections make direct excitatory connections in contralateral CA1

Does a relatively small number of CA3 neurons targeted in SST-Cre mice have a measurable functional impact on neuronal networks? CA3 pyramidal neurons are known to make extensive long-range connections to the contralateral hippocampus (Buzsáki & Czéh, 1981; Buzsáki & Eidelberg, 1982; Finnerty & Jefferys, 1993) and the septum (Risold & Swanson, 1997). We therefore examined if the small number of CA3 neurons targeted in SST-Cre mice is sufficient to generate detectable contralateral projections. Unilateral rAAV injection in the CA3 region of SST-Cre mice led to a strong axonal EYFP signal in the contralateral hippocampus (**Figure 2A**) and the septum (**Figure 2B**). The axon distribution was as described for CA3 pyramidal cells, with EYFP-expressing axons mainly in SO and SR of both the CA1 and CA3 regions.

Contralateral projections have been described not only for CA3 pyramidal neurons, but also for inhibitory hippocampal interneurons including SST-expressing subtypes (Eyre & Bartos, 2019; Zappone & Sloviter, 2001). We therefore went on to further characterize the functional properties of contralaterally projecting axons, with the goal to assess i) if they correspond to excitatory projections arising from CA3 pyramidal neurons and ii) if they are sufficiently numerous to cause significant physiological effects. To this end, we obtained patch-clamp recordings from CA1 pyramidal neurons in mice expressing hChR2 in the contralateral CA3 region in SST-Cre mice. This allowed us to perform light-based stimulation of contralaterally projecting axons while recording from CA1 pyramidal neurons. To separate excitatory from inhibitory neurotransmission, we voltage clamped CA1 neurons to different holding voltages. Currents at −80 or −70 mV were evoked close to the chloride reversal potential and are therefore dominated by excitatory postsynaptic currents (EPSCs), whereas currents evoked at 0 or −10 mV are dominated by inhibitory postsynaptic currents (IPSCs). In all CA1 pyramidal neurons, blue light illumination reliably evoked both excitatory and inhibitory currents (**Figure 2C,D,E**). To ascertain which of these components are monosynaptic in nature, we applied the Na^+^ channel blocker tetrodotoxin (TTX, 1µM), which invariably blocked synaptic transmission completely. Coapplying TTX with 4-aminopyridine (4-AP, 200µM) enables direct light-based transmitter release from terminals expressing ChR2, and thus indicates monosynaptic connections. Coapplication of 4-AP recovered EPSCs, but not IPSCs (**Figure 2C**; EPSCs 217%, IPSCs 1% of baseline). The recovery of EPSCs but not IPSCs indicates that contralateral projections in SST-Cre mice are excitatory. Additionally, these results indicate that the light-evoked IPSCs are due to polysynaptic recruitment of interneurons. This idea is supported by the temporal delay between excitatory and inhibitory conductances (**Figure 3G**). Consistent with polysynaptic recruitment of inhibitory interneurons, light-evoked IPSCs were abrogated by blocking glutamatergic transmission with CNQX (50µM) and D-AP5 (200µM; **Figure 2D;** EPSCs 29%, IPSCs 8% of baseline). Finally, we show that – as expected – light-evoked IPSCs were sensitive to the GABA-A blocker gabazine (10µM; **Figure 2E**; EPSCs 114%, IPSCs 14% of baseline).

**Figure 3:**
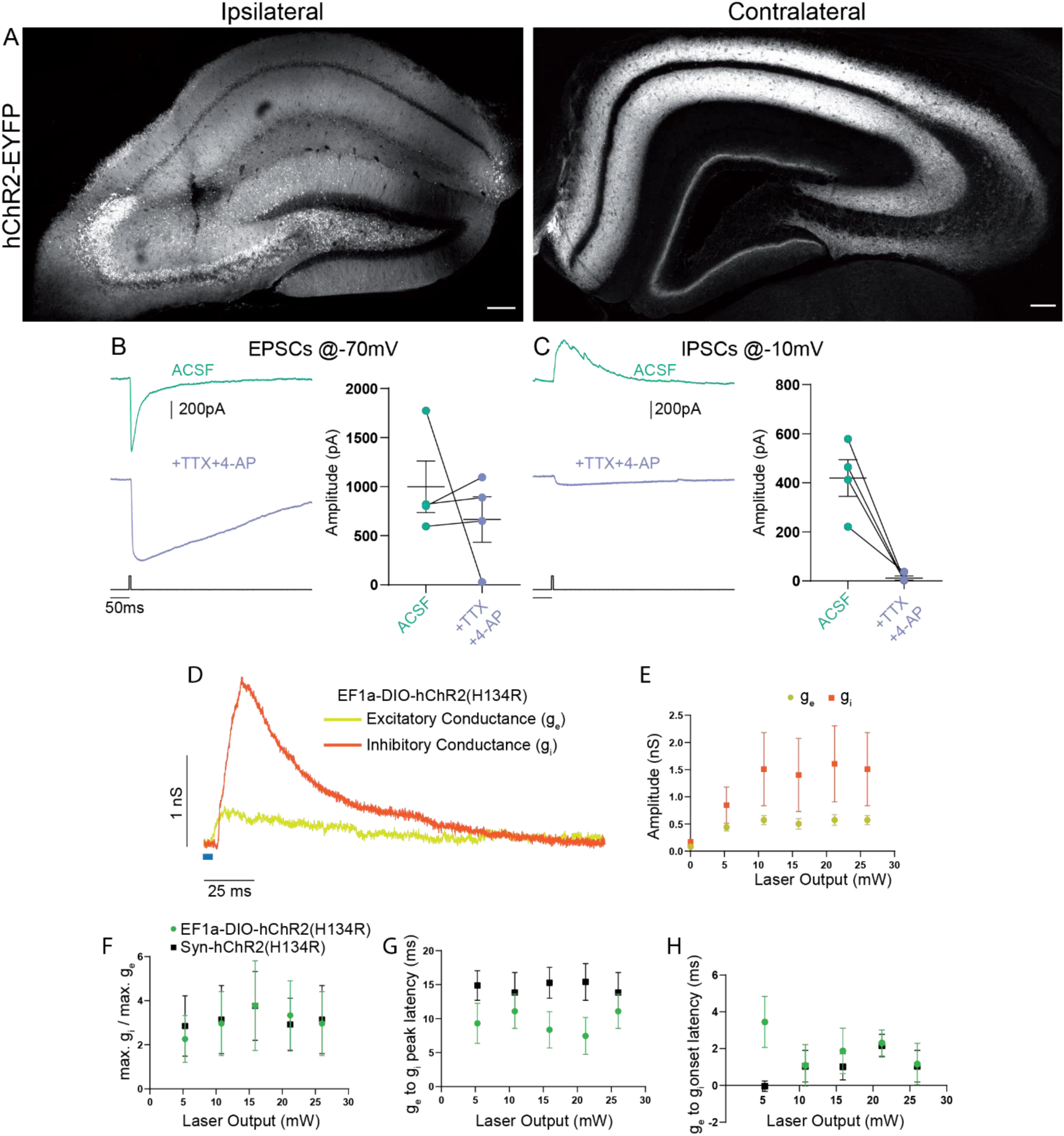
Contralateral projections originating from Cre-expressing cells in CA3 vs the general CA3 neuron population are functionally indistinguishable. A) Confocal images from post-fixed acute slices of the ipsilateral injection site (left) and the contralateral hippocampus (right). Unconditional viral expression. B) EPSCs and C) IPSCs (right) before and after bath application of TTX and 4-AP measured in contralateral CA1 pyramidal cells. 5ms light stimulation at 26mW fiber output. Ratio t-test of dependent samples one-tailed: EPSCs, p= 0,2284, t=0,8519, df=3; IPSCs, p= 0,0069, t=5,200, df=3. D) Example from conductance analysis of fibers in the SST-Cre mouse line, conditionally expressing. 26mW light fiber output and 5 ms light stimulation. Excitatory conductance was calculated from gabazine traces. Inhibitory conductance was calculated from gabazine subtracted traces. E) Quantification of excitatory and inhibitory peak conductance at different laser powers. 2-way ANOVA Greenhouse-Geisser corrected: Main effects, Laser Output: p=0.0422, DF = 5, F(1.182, 7.091) = 5.849, Conductance Type,: p=0.2189, DF=1, F(1.000, 6.000)=1.885. Interaction: p=0.2527, DF=5, F(1.115, 6.693)=1.600. F) Quantification of conductance ratios (inhibitory peak conductance divided by excitatory peak conductance) for conditional viral expression (EF1a-DIO-hChR2(H134R)) and unconditional expression (Syn-hChR2(H134R)). 2-way ANOVA Greenhouse-Geisser corrected: Main effects, Laser Output: p=0.1406, DF = 4, F(1.393, 15.33)=2.341, Expression Type: p=0.9614, DF=1, F(1, 11)=0.002455. Interaction: p=0.7974, DF=4, F(4, 44)=0.4143. G) Quantification of latency between excitatory peak conductance and inhibitory peak conductance. 2-way ANOVA Greenhouse-Geisser corrected: Main effects: Laser Output: p=0.6446, DF=4, F(1.720, 18.92)= 0.4014, Expression Type: p=0.1766, DF=1, F(1, 11)=2.085. Interaction: p=0.0320, DF=4, F(4, 44)=2.912. H) Quantification of latency between excitatory conductance onset and inhibitory conductance onset. 2-way ANOVA Greenhouse-Geisser corrected: Main effects, Laser Output: p=0.6474, DF=4, F(2.306, 25.37)=0.4853, Expression Type: p=0.1759, DF=1, F(1,11)=2.092. Interaction: p=0.3588, DF=4, F(4,44)=1.121.

Taken together, we found no evidence for direct commissural inhibition from SST^+^ interneurons from CA3 to CA1. Instead, direct excitatory transmission recruited strong polysynaptic inhibition.

### 4.3 Unconditionally transduced and SST-Cre fibers are functionally indistinguishable in Contralateral CA1

To investigate if this is consistent with the canonical CA3 to CA1 commissural projection, we induced broad expression of ChR2 in all CA3 cell types using viral gene transfer of an unconditional construct leading to expression of EYFP-hChR2. Light based manipulations should be dominated by activity of pyramidal cells, since they vastly outnumber other neuronal subtypes. Virus injection resulted in strong fluorescence signal in CA1, CA3 and DG that was dominated by fiber signal at the injection site (**Figure 3A**). Contralateral to the injection site, we found prominent labelling of axons in CA1 and C3 in both SR and SO as well as the inner molecular layer of the DG. The DG fiber pattern was consistent with the commissural mossy cell projection and the fiber patterns in CA1 and CA3 with the commissural CA3 projection. We again assessed the monosynaptic transmission onto contralateral CA1 pyramidal cells using combined application of TTX and 4-AP (1µM, 200µM) and found that it completely inhibited IPSCs (**Figure 3C;** EPSCs 88%, IPSCs 4% of baseline). Next, we asked if there are quantitative differences between the SST-Cre fibers and the unconditionally transduced fibers. We converted the pharmacologically isolated currents (**Figure 2E**) to conductances (**Figure 3D**) according to holding and reversal potentials (see Methods). Because the density of EYFP-hChR2 positive fibers is much larger in the unconditional case, the absolute conductances cannot be compared meaningfully. However, because the inhibition is polysynaptic, it is expected to scale to some extent with the excitation. Therefore, the ratio between excitation and inhibition can give insights into differential recruitment in the micronetwork.

We found that in the SST-Cre line, the inhibitory conductance was stronger than the excitatory one (**Figure 3D,E**). Comparing the SST-Cre line with the unconditional case, we did not detect a difference between the ratios of maximum inhibition and excitation (**Figure 3F**). In both cases, the amplitude of inhibition was larger than that of inhibition for different strengths of light-based stimulation. Furthermore, the latencies between the onset of excitation and inhibition showed no significant difference (**Figure 3H**) and were consistent with values found in CA3 to CA1 Schaffer collateral projections (Scanziani, 2001). However, the latencies between the peak of the excitatory conductance and the inhibitory conductance showed a significant interaction between laser output and the type of expression. Main effects were not significant (**Figure3G**, Greenhouse-Geisser corrected 2-way ANOVA).

### 4.4 Commissural CA3 fibers make synaptic contacts on spines and originate primarily from PCL cells

To further confirm that contralateral projections are excitatory, we used miniSOG photooxidation to generate electron-dense labelling in contralateral CA1 SO localized to fibers with Cre recombinase activity in the SST-Cre line (**Figure 4**). Of 70 miniSOG positive structures, 40 clearly were presynaptic boutons making postsynaptic contacts. All 40 structures made contact on a spine, 4 of them made contact on 2 spines (**Figure 4A**). Serial imaging sections of 25 boutons showed that 22 of them unambiguously made contact on spines (**Figure 4B**). The other three boutons were not entirely sectioned. The types of most synaptic contacts could not be defined clearly because of the electron dense labelling in the pre-synapse. However, the postsynaptic densities that are clearly in the imaging plane appear asymmetric. Together with the fact that they all contact spines, this data suggest that the direct contacts are predominantly excitatory, and we found no evidence for direct inhibitory contacts in CA1 SO.

**Figure 4:**
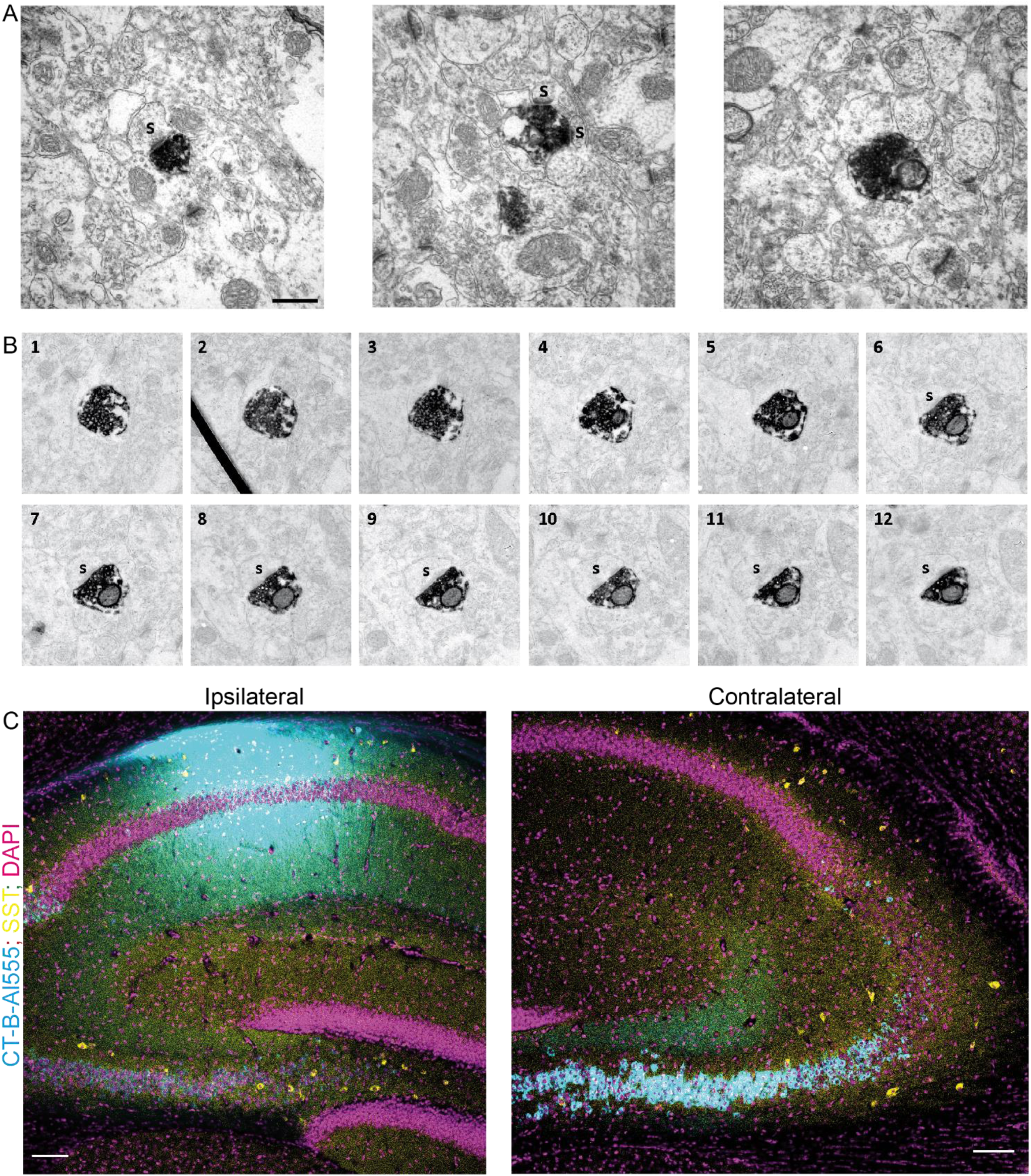
Contralaterally projecting axons originating from Cre-expressing neurons in CA3 are excitatory. A) miniSOG positive electron dense structure making presynaptic contact on one spine (left) two spines (middle) and no identifiable contact (right). Scalebar 500nm. B) Serial scan of a miniSOG positive electron dense structure. Post synaptic spine is first identifiable in slice 6. C) Choleratoxin-B tracing in CA1. Ipsilateral injection of CT-B subunit in CA1. Contralateral, retrogradely traced cells (cyan) and SST immunoreactive cells (yellow). Scalebar 200µm.

Next, we used retrograde tracing in CA1 with CT-B to determine which cell types project to contralateral CA1 (**Figure 4C**). We found that virtually all projecting cells were in the CA3 pyramidal cell layer. With the SST staining we identified 81 cells, none of which was CT-B positive. This data suggests that somatostatin interneurons are not part of the commissural projection.

Finally, we tried to relate our findings to other commonly used Cre mouse lines. Therefore, we used data from the Allen Brain Institute to estimate specificity in two other interneuron targeting Cre lines.

### 4.5 Unspecificity is comparable in another SST-Cre mouse line

The mouse line we describe here has not been widely used. The SST-IRES-Cre line (Taniguchi et al., 2011) is much more wide-spread, with 203 publications relating to it according to Jackson Laboratories (as of 11.10.2019). To relate our findings to the SST-IRES-Cre line we used the Allen Brain Institute’s transgenic characterization data of the mouse connectome project (Oh et al., 2014). We used two experiments in which the SST-IRES-Cre mouse line was crossed with a tdTomato expressing reporter line (Ai-14) and fluorescent in-situ hybridization (FISH) was performed for SST.

We found that the SST-IRES-Cre mouse line is similarly unspecific in CA3, with only 48/127 (37.8%) tdTomato^+^ cells being SST-mRNA^+^ in the PCL, 82/100 (82%) in SO and 61/74 (82.4%) in SR (**Figure 5B,C**). The CA1 area also contained some SST-cells in the PCL but appeared overall more specific with 29/51 (56.9%) tdTomato^+^ cells being SST-mRNA positive, 281/299 (94%) in SO and 20/24 (83.3%) in SR (**Figure 5D,E**). The total amount of cells located in PCL is much smaller in the SST-IRES-Cre line (**Figure 5A**) than the SST-Cre line when crossed with the Ai14 reporter line (**Figure 1C**).

**Figure 5:**
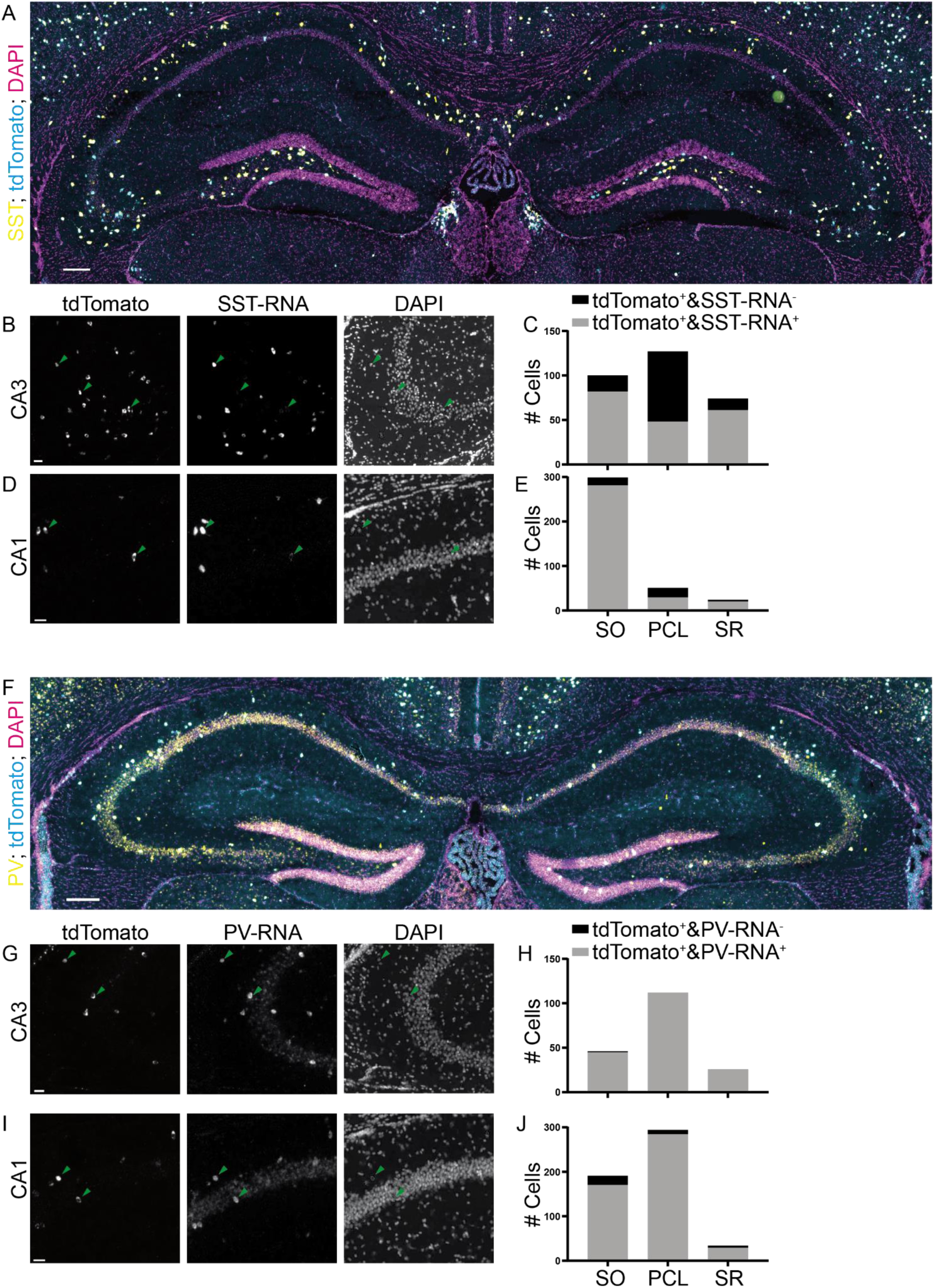
Analysis of Cre lines. Images from the Allen Brain Institute. Cre mouse lines were crossed with the tdTomato reporter line Ai14. A) Experiment 167643437, image ID 167643516. Contrast auto adjusted and lookup tables change. B-E) Example images cropped from A), contrast unadjusted. Quantification on the right. F) Experiment 111192541, image ID 111192610. Contrast auto adjusted and lookup tables changed. G-J) Example images cropped from F) contrast unadjusted. Quantification on the right. Scalebars: 100 and 20µm.

We also quantified colocalization of Cre-induced recombination with PV expression in the Pvalb-IRES-Cre mouse line (Hippenmeyer et al., 2005). We found that this mouse line was much more specific than both SST-Cre mouse lines in both the CA3 and CA1 regions (**Figure 5F-J**; CA3: 45/46, 97.8% SO; 112/112, 100% PCL; 26/26, 100% SR. CA1: 170/191, 89% SO; 284/294, 96.6% PCL; 29/34, 85.3% SR). In summary, the problems we described are common to at least one other commonly used SST-Cre line, but less severe in another mouse line used to study interneurons.

## 5 Discussion

We show for the first time that CA3 PCs that are unspecifically targeted in an SST-Cre mouse line make functional connections indistinguishable from those of canonical CA3 PCs. While the specificity of SST-Cre lines has been questioned before, the functional relevance of unspecific expression of Cre-recombinase was unknown. Estimating the potential effects of unspecific expression is essential for neuronal perturbation studies that seek to isolate the function of specific cell-types. Our data suggest that studies that perturb SST cells in CA3 with the two examined SST-Cre lines would be massively confounded by Cre recombinase expression in CA3 pyramidal cells.

### 5.1 How relevant are these findings for other Cre mouse lines?

We demonstrate wide-spread physiological effects of unspecific Cre-expression in only one mouse line, but have identified unspecific expression in another, widely used mouse line following quantitative analysis of Allen Brain Atlas data. Indeed, specificity issues with an SST-Cre mouse line were raised previously (Taniguchi et al., 2011). Moreover, a further study has found targeting of a large number (31%) of slow-spiking cells in the CA1 PCL, also consistent with unspecific expression (Mikulovic et al., 2015). It is worth noting that the mouse line we used was generated by the same method as most modern Cre mouse lines, the BAC technology. Specificity can vary widely between Cre lines and brain areas, as our comparison of the SST-IRES-Cre and the Pvalb-IRES-Cre lines shows. Therefore, specificity should not be generalized lightly to other Cre mouse lines or even to other brain areas in the same mouse line. We suggest that pending careful quantitative analysis in all the subregions under investigation in the specific study, caution is warranted in assuming specificity.

### 5.2 Do SST-expressing interneurons make contralateral connections?

Despite the specificity issues, the SST-Cre mouse line clearly targets SST^+^ INs in CA3. We found no evidence for direct inhibition from those cells onto contralateral CA1 PCs in our patch clamp experiments. Even the unconditionally transduced slices did not reveal monosynaptic inhibition, despite targeting all inhibitory cell types. Furthermore, our anatomical EM data showed no evidence for inhibitory synapses in contralateral CA1 SO. The CT-B data did not reveal cells outside CA3 PCL projecting to contralateral CA1. This leads us to the conclusion that an inhibitory CA3 to contralateral CA1 connection is nonexistent or extremely weak and SST^+^ interneurons do not contribute to it.

Although we focused on CA3 and CA1, we noted very sparse fiber signal in the outer molecular layer of DG in the SST-Cre line. This is in line with previous anatomical evidence showing a commissural projection with a GABAergic component (Deller, Nitsch, & Frotscher, 1995; Zappone & Sloviter, 2001). However, using in-vivo patch clamp and optogenetics we did not find evidence for a functional connection onto granule cells (data not shown).

Recently, Eyre & Bartos (2019) found commissural DG fibers in a GAD2-Cre and the SST-Cre mouse line we used in our study. They performed whole-cell patch clamp in contralateral GCs and found no evidence for a functional connection, in line with our findings. However, they also report commissural fibers from CA3 to CA1 in both mouse lines. Our findings suggest that the contralateral CA1 fiber signal originates from CA3 pyramidal cells, rather than SST INs. Furthermore, our findings do not explain the CA1 fiber labeling in the GAD2-Cre line, but they suggest that the functional aspect of this inhibitory connection is weak. It remains to be shown which inhibitory cell type, if any, is responsible for this contralateral fiber signal.

### 5.3 Utility of mouse lines with unspecific principal cell expression for in-vivo experiments

A common use of Cre lines is circuit perturbation during behavioral tasks. Principal cell connections can span wide areas of the brain and must be accounted for when studying interneurons. When light is delivered to the brain through light fibers, it can travel considerable distances. Therefore, light delivered to areas where transgene expression is specific, could affect unspecifically expressing cells and fibers in faraway areas. Notably, such effects cannot be excluded with a commonly used control group expressing only GFP (or another fluorophore) instead of a light sensitive opsin. The same applies to a larger extent to chemogenetic experiments, where the agonist might be delivered systemically, rather than locally.

To ensure that principal cell expression does not confound a behavioral experiment, the colocalization between transgene expressing cells and the appropriate interneuron marker should be quantified for all areas where viral transduction occurred. This includes the injection cannula tract. When the transgene is expressed by crossing mouse lines, the expressing fiber distribution throughout the entire brain should be examined carefully. Especially for optogenetic experiments, it would be valuable to additionally check for direct excitatory synaptic transmission. For a specific mouse line, no direct excitatory currents should be detectable. Importantly, the net effect of a direct excitatory connection can be reduced spiking through recruitment of feedforward and feedback inhibition (Buzsáki & Czéh, 1981). Therefore, it is not sufficient to quantify spiking or activity levels in the post-synaptic population to exclude direct excitation. These issues should be taken into account when using any Cre-mouse line for in-vivo behavioral experiments, particularly the SST-Cre mouse lines used in the present study.

## 6 Acknowledgement

We acknowledge the support of the Microscopy Core facility of the University Bonn Medical Center, specifically Hannes Beckert, and the Viral Core Facility, specifically Susanne Schoch-McGovern. We thank Joanna Komorowska-Müller for helpful suggestions on the antibody staining protocol and figure design. We also thank Kristina Piwellek for excellent technical assistance with viral injections for miniSOG labeling.

## 8 Data Availability Statement

Datasets are available on request. The raw data supporting the conclusions of this manuscript will be made available by the authors, without undue reservation, to any qualified researcher.

## 10 Author Contributions

DMK and HB designed the project and wrote the manuscript. DMK performed viral injections, SST antibody stainings, patch clamp recordings and quantification of Allen Brain Institute Data. TO performed viral injections for miniSOG and performed photooxidation. MS acquired EM images and quantified them. SE performed CT-B injections and SST antibody stainings in connection with them.

